# Aberrant enteric neuromuscular system and dysbiosis in amyotrophic lateral sclerosis

**DOI:** 10.1101/2021.07.13.452097

**Authors:** Yongguo Zhang, Destiny Ogbu, Shari Garrett, Yinglin Xia, Jun Sun

## Abstract

**Background:** Emerging evidence has demonstrated that microbiota directly affects the enteric neuron system (ENS) and smooth muscle cell functions via metabolic products or endogenous bacterial components. Amyotrophic Lateral Sclerosis is a neuromuscular disease characterized by the progressive death of motor neurons and muscle atrophy. The GI symptoms in patients were largely ignored or underestimated, especially before the diagnosis of ALS. The relationship between enteric neuromuscular system and microbiome in ALS progression is unknown.

**Methods:** We performed longitudinal studies on the ENS and microbiome in the ALS human-SOD1^G93A^ transgenic G93A mice. We treated age-matched wild-type and ALS mice with bacterial product butyrate or antibiotics to investigate microbiome and neuromuscular functions. Intestinal motility, microbiome, an ENS marker GFAP, a smooth muscle marker (SMMHC), and human colonoids have been examined. The distribution of human-G93A-SOD1 (Superoxide Dismutase 1) protein was tested as an indicator of ALS progression.

**Results:** At 2-month-old before ALS onset, G93A mice had significant lower intestinal motility, decreased grip strength, and reduced time in the rotarod. We observed increased GFAP and decreased SMMHC expression. These changes correlated with consistent increased aggregation of mutated SOD1^G93A^ in the colon, small intestine, and spinal cord. Butyrate and antibiotic treatment showed a significantly longer latency to fall in the rotarod test, reduced SOD1^G93A^ aggregation, and enhanced ENS and muscle function. Feces from 2-month-old SOD1^G93A^ mice significantly enhanced SOD1^G93A^ aggregation in human colonoids transfected with a SOD1^G93A^-GFP plasmid. Longitudinal studies of microbiome data further showed the altered bacterial community related with autoimmunity *(e.g., Clostridium sp. ASF502, Lachnospiraceae bacterium A4)*, inflammation (e.g., *Enterohabdus Muris*,), and metabolism (e.g., *Desulfovibrio fairfieldensis*) at 1- and 2-month-old SOD1^G93A^ mice, suggesting the early microbial contribution to the pathological changes.

**Conclusions:** We have demonstrated a novel link between microbiome, hSOD1^G93A^ aggregation, and intestinal mobility. Dysbiosis occurred at the early stage of the ALS mice before observed mutated-SOD1 aggregation, slow intestinal motility, and dysfunction of ENS. Manipulating the microbiome improves the muscle performance of SOD1^G93A^ mice. Our study provides insights into fundamentals of intestinal neuromuscular structure/function and microbiome in ALS.

## Introduction

ALS is characterized by the progressive death of motor neurons and muscle atrophy. Early diagnosis of ALS has been a long-standing challenge in the field. There are case reports that celiac disease with neurologic manifestations was misdiagnosed as ALS ^1–3^. Researchers in Israel discovered a possible link between ALS and sensitivity to gluten ^4^. In certain cases, an ALS syndrome might be associated with autoimmunity and gluten sensitivity. These reports indicate the GI symptom in ALS and challenge in diagnosis of diseases. Although the data are preliminary, gluten sensitivity is potentially treatable; the diagnostic challenge should not be overlooked. Moreover, inflammatory cytokines (e.g., IL-6) and bacterial product LPS were elevated in ALS ^5, 6^. Autoimmune disease (e.g., Crohn’s disease) associations with ALS raise the possibility of shared genetic or environmental risk factors in the pathogenesis and progression of ALS ^7^.

Altered intestinal homeostasis and microbiome contribute to a variety of neurological diseases (e.g. autism, Alzheimer’s disease, and Parkinson’s disease). We are the first to report the elevated intestinal inflammation, reduced beneficial bacteria, and shift of microbiome profile in ALS ^8, 9^ ^10^. Later, there are studies from several groups reporting the dysbiosis in human ALS and experimental animal models ^11–14^. Thus, the current evidence indicates that intestinal dysfunction and dysbiosis may actively contribute to ALS pathogenesis.

Intestinal motility is a key physiologic parameter governing digestion and absorption of nutrients affected by ENS ^15^, microbiome, and host genetics ^16–20, 18, 21^. The ENS ^15^ is an important regulator of the proliferation and differentiation of the mucosal epithelium; serotonergic neurons in the myenteric plexus activate submucosal cholinergic neurons that innervate intestinal crypts to stimulate proliferation, thus playing an essential role in various neurodegenerative diseases ^22, 23^ ^24^. However, studies of the ENS, motility, and microbiome in ALS are lacking.

In the current study, we examined the changes in enteric neurons and smooth muscle in the prodromal phase of ALS, using an ALS human-SOD^G93A^ transgenic G93A mice, with or without butyrate treatment. We chose human-SOD^G93A^ transgenic mice because a fraction of familial ALS is associated with mutations in the superoxide dismutase gene (SOD1) ^25^. Mouse models expressing ALS-linked human-SOD1 mutations effectively recapitulate many features of the human disease and have been extensively used to investigate pathogenic mechanisms of ALS ^26^. We performed longitudinal studies on the ENS, microbiome, and SOD1 aggregation in the ALS G93A mice. Human colonoids transfected with SOD1^G93A^-GFP plasmids were used for the microbial-host interactions and SOD1 aggregation. We investigated the mechanism in altered ENS and intestinal motility that contribute to ALS progression, correlated with SOD1^G93A^ aggregation, and dysbiosis. Our data suggest that modulation microbiome by treating the mice with beneficial butyrate or antibiotics in ALS changes the ENS and disease progression. This study provides insights into fundamentals of intestinal neuromuscular structure/function and microbiome in ALS. Better understanding the intestinal dysfunction and dysbiosis in ALS will help for the early diagnosis and development of new treatments.

## Results

### Slow intestinal motility and weak muscle strengths correlated with aggregation of SOD1^G93A^ in intestine of the ALS mice

To study the intestinal changes before the onset of ALS symptoms, we assessed the intestinal transit time of G93A mice at 1-, 2-, and 3-month-old, using a whole gut motility assay ^27^. The 1-month-old G93A mice showed no significant changes of motility, compared with the wild-type (WT) mice. We found significantly increased gut transit time starting at 2-month-old (Fig. 1A), suggesting slow intestinal motility in the G93A mice. The disturbance of motility in ALS may reflect changes in smooth muscle function. We then examined the neuromuscular activity performance o*f* the G93A mice on an accelerating rotarod ^28^. Muscle strength was evaluated using the grip strength and the hanging wire test. We then tested their ability of riding rotarod. Starting at 2-month-old, G93A mice had significantly reduced time in the rotarod, an indicator of weakened muscle function (Fig. 1B). We further examined the forelimb grip strength (Fig. 1C). Interestingly, the changes of intestinal motility were negatively correlated with the weakened muscles in the grip strength (r=-0.68, p=0.015 in month 2 and r=-0.83, p<0.0001 in month 3, respectively). We also found the similar changes of hindlimb grip strengths (Fig. 1D) (r=-0.48, p=0.1172 in month 2 and r=-0.83, p<0.0001 in month 3, respectively).

**Figure 1.**
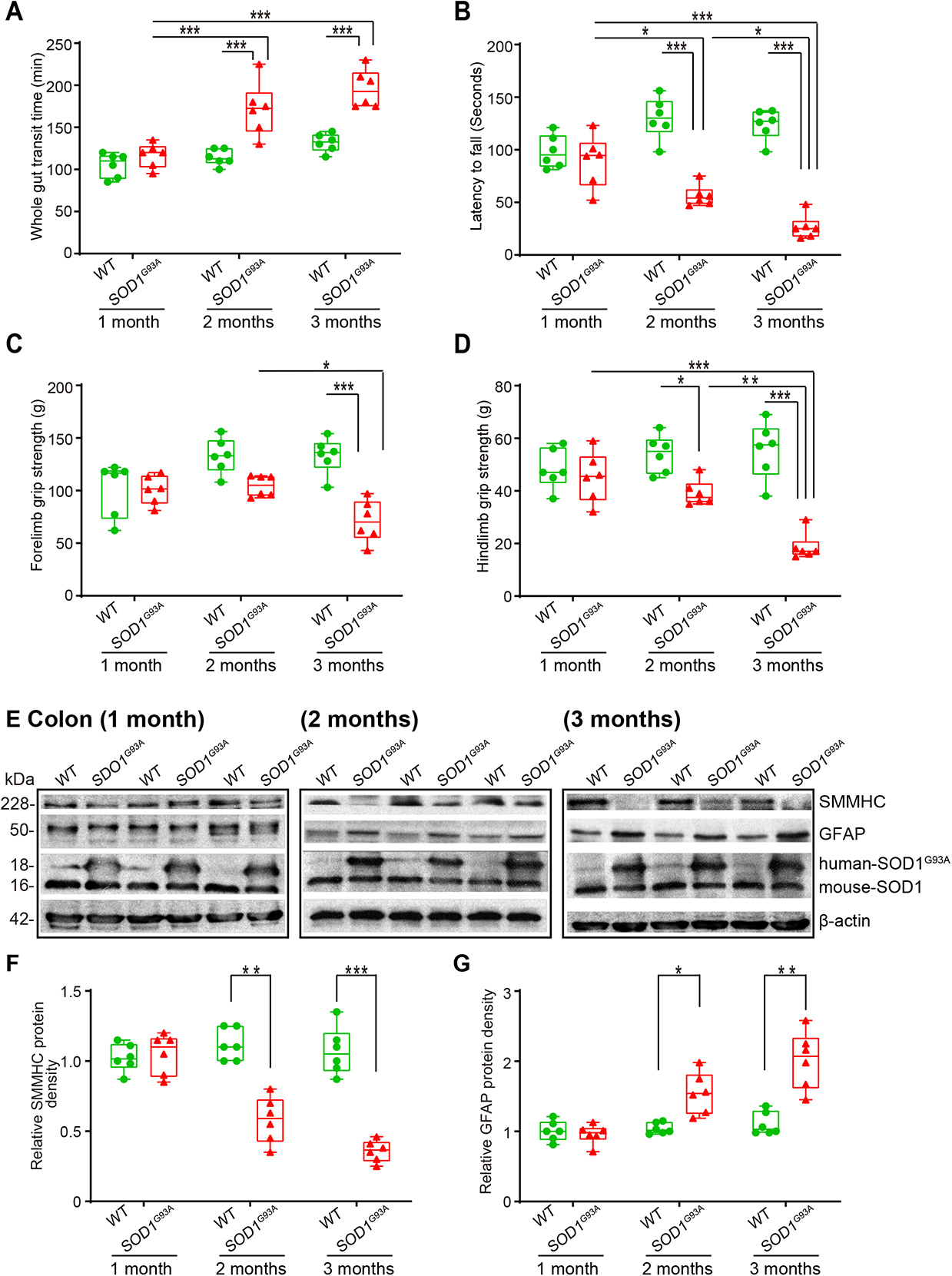
SOD1^G93A^ mice have slow intestinal motility, decreased rotarod test time and grip strength during ALS progression. ***(A)*** SOD1^G93A^ mice significantly increased gut transit time starting at 2-month-old compared to WT mice. Age-matched WT mice and SOD1^G93A^ mice were tested in intestinal motility using Evans blue marker (5% Evans blue, 5% gum Arabic in drinking water). (Data are expressed as mean ± SD. n = 6, two-way ANOVA test, ***P < 0.001, adjusted by the Tukey method). ***(B)*** Starting at 2-month-old, SOD1^G93A^ mice had significant reduced rotarod test time compared to WT mice. (Data are expressed as mean ± SD. n = 6, two-way ANOVA test, *P < 0.05, **P < 0.01, ***P < 0.001, adjusted by the Tukey method). ***(C)*** Forelimb grip strength in WT and SOD1^G93A^ mice at different time points and ***(D)*** Hindlimb grip strength in WT and SOD1^G93A^ mice at different time points. (Data are expressed as mean ± SD. n = 6, two-way ANOVA test, *P < 0.05, **P < 0.01, ***P < 0.001, adjusted by the Tukey method). ***(E)*** At the age of 2 months old, the expression of SMMHC protein started to decrease while the expression of GFAP protein started to increase in age matched SOD1^G93A^ mice compared to WT mice, and ***(F)*** and ***(G)*** Quantification for the expression of SMMHC and GFAP proteins in different time points. (Data are expressed as mean ± SD. n = 6, Kruskal-Wallis test with pairwise comparisons using Wilcoxon rank sum exact test, *P < 0.05, ***P < 0.01, ***P < 0.001 adjusted by the FDR method).

### Altered enteric neuromuscular markers in the ALS G93A mice

At the protein levels, we then examined the expression of the smooth muscle marker, smooth muscle myosin heavy chain (SMMHC) ^29^, using western blots (WB) (Fig. 1E). There was a significant reduction of SMMHC at the protein level in the 2-month-old G93A mice (male and female) (Fig. 1E **&** 1F). There are enteric glia cells in the ENS and beneath the epithelium throughout the intestinal mucosa ^30^. A recent study showed that enteric glia cells regulate intestinal motility ^31^. Therefore, we investigate the enteric glia by testing glial fibrillary acidic protein (GFAP) in G93A mice at the ages of 1-, 2-, and 3-month-old (Fig. 1E), whereas the endogenous mouse SOD remained the same. The GFAP expression was significantly enhanced in the 2-month-old and 3-month-old G93A mice (Fig. 1F **&**1G). We also showed a significant increase of human SOD1^G93A^ in the intestine in the 2-month-old transgenic G93A mice, and the changes of ENS markers were correlated with the increased human SOD1G93A in the colon in the G93A mice (Fig. 1E).

### Altered enteric neuromuscular structure in the ALS G93A mice starting at 2-month-old

Using immunostaining, we further detected the distribution of SMMHC in the intestine of G93A mice in different age groups (Fig. 2A). Starting from 2-month-old, we were able to find the reduced SMMHC in the colon of the G93A mice. In the meantime, there is no change of alpha-smooth muscle actin in the colon of the G93A mice (Fig. 2A). In the 2-month-old G93A mice, we found enhanced GFAP (Fig. 2B). The intracellular neuronal marker protein gene product (PGP) 9.5 is a label for neural-crest-derived precursor cells during gut development ^32^. PGP9.5 was also enhanced in the G93A mice (Fig. 2C). The distribution of human-G93A-SOD1 mutated protein was used as an indicator of ALS progression. Interestingly, we observed the aggregation of SOD1^G93A^ in the intestine, starting at the 2-month-old (Fig. 2D).

**Figure 2.**
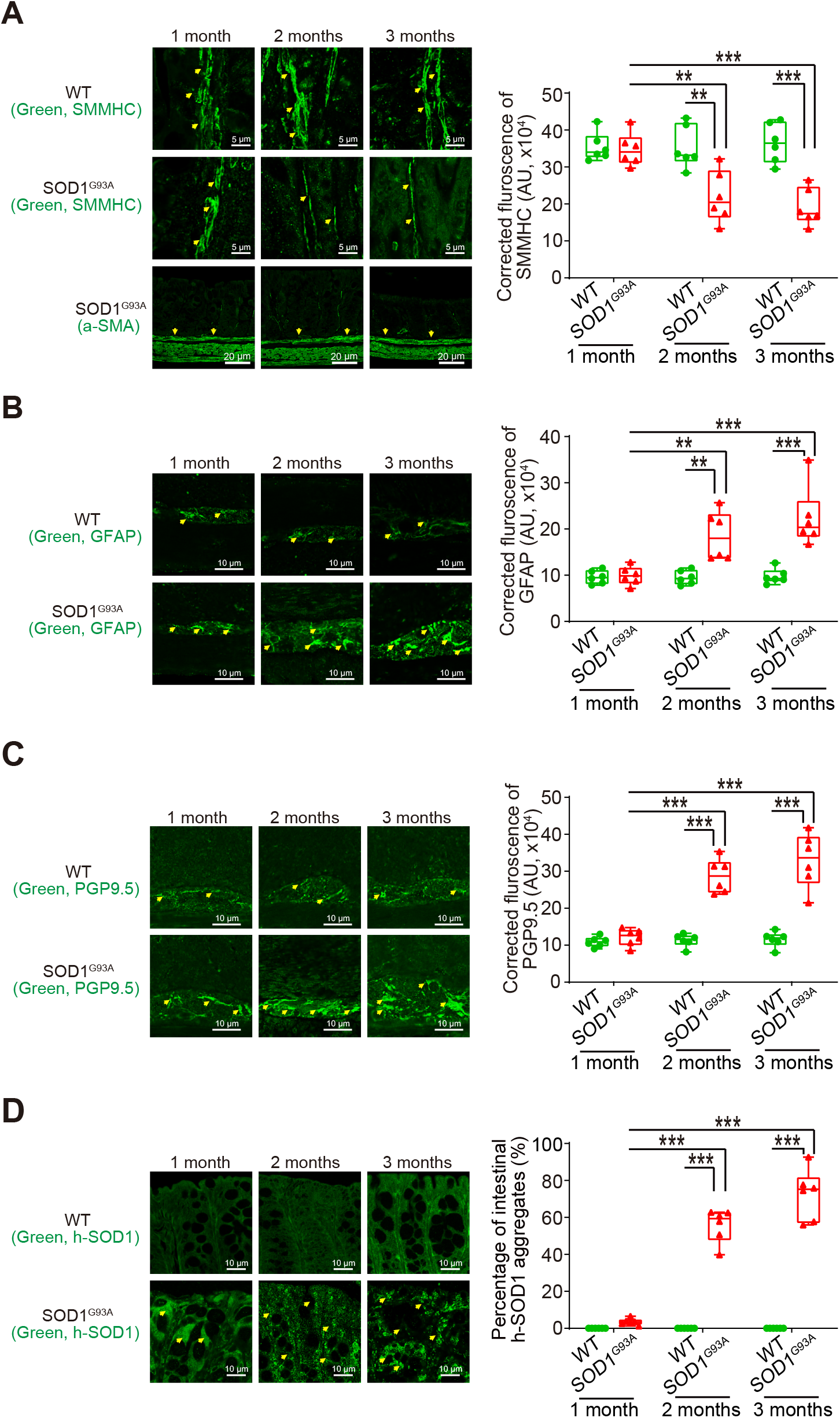
Defects in the enteric nervous system structure in the SOD1^G93A^ mice starting at 2-month-old. ***(A)*** SMMHC started decrease at 2-month-old in the colon of the SOD1^G93A^ mice compared to the aged-match WT mice in immunofluorescence staining and quantification of SMMHC staining. There is no change of alpha-smooth muscle actin in immunofluorescence staining. Images are from a single experiment representative of 6 mice per group. (Data are expressed as mean ± SD. n = 6, two-way ANOVA test, **P < 0.01, ***P < 0.001, adjusted by the Tukey method). ***(B)*** GFAP started to increase at 2-month-old in the colon of the SOD1^G93A^ mice compared to the age-matched WT mice in immunofluorescence staining and quantification of GFAP staining. Images are from a single experiment, representative of 6 mice per group. (Data are expressed as mean ± SD. n = 6, Kruskal-Wallis test, ***P < 0.01, ***P < 0.001, adjusted by the FDR method). ***(C)*** PGP9.5 was also enhanced in the SOD1^G93A^ mice at 2-month-old in the colon of the SOD1^G93A^ mice compared to the age-matched WT mice in immunofluorescence staining and quantification of PGP9.5 staining. Images are from a single experiment, representative of 6 mice per group. (Data are expressed as mean ± SD. n = 6, two-way ANOVA test, ***P < 0.001). ***(D)*** Aggregation of human-SOD1^G93A^ protein was observed starting from 2-month-old SOD1^G93A^ mice compared to the age-matched WT mice in immunofluorescence staining and human-SOD1 protein aggregates percentage analysis. Images are from a single experiment, representative of 6 mice per group. (Data are expressed as mean ± SD. n = 6, Kruskal-Wallis test with pairwise comparisons using Wilcoxon rank sum exact test, **P < 0.01, ***P < 0.001, adjusted by the FDR method).

### The aggregation of SOD1^G93A^ in the neurons and intestine of G93A mice

The aggregation of SOD1^G93A^ in the spinal cord and neurons is a molecular hallmark of ALS progression. The aggregation of human-SOD1^G93A^ protein was observed in the lumbar spine white matter and gray matter, starting in 2-month-old SOD1^G93A^ mice (Fig. 3A **&** B). GFAP expression was also enhanced starting from 2-month-old in the white matter and gray matter of the lumbar spine in the SOD1^G93A^ mice (Fig. 3C **&** D).

**Figure 3.**
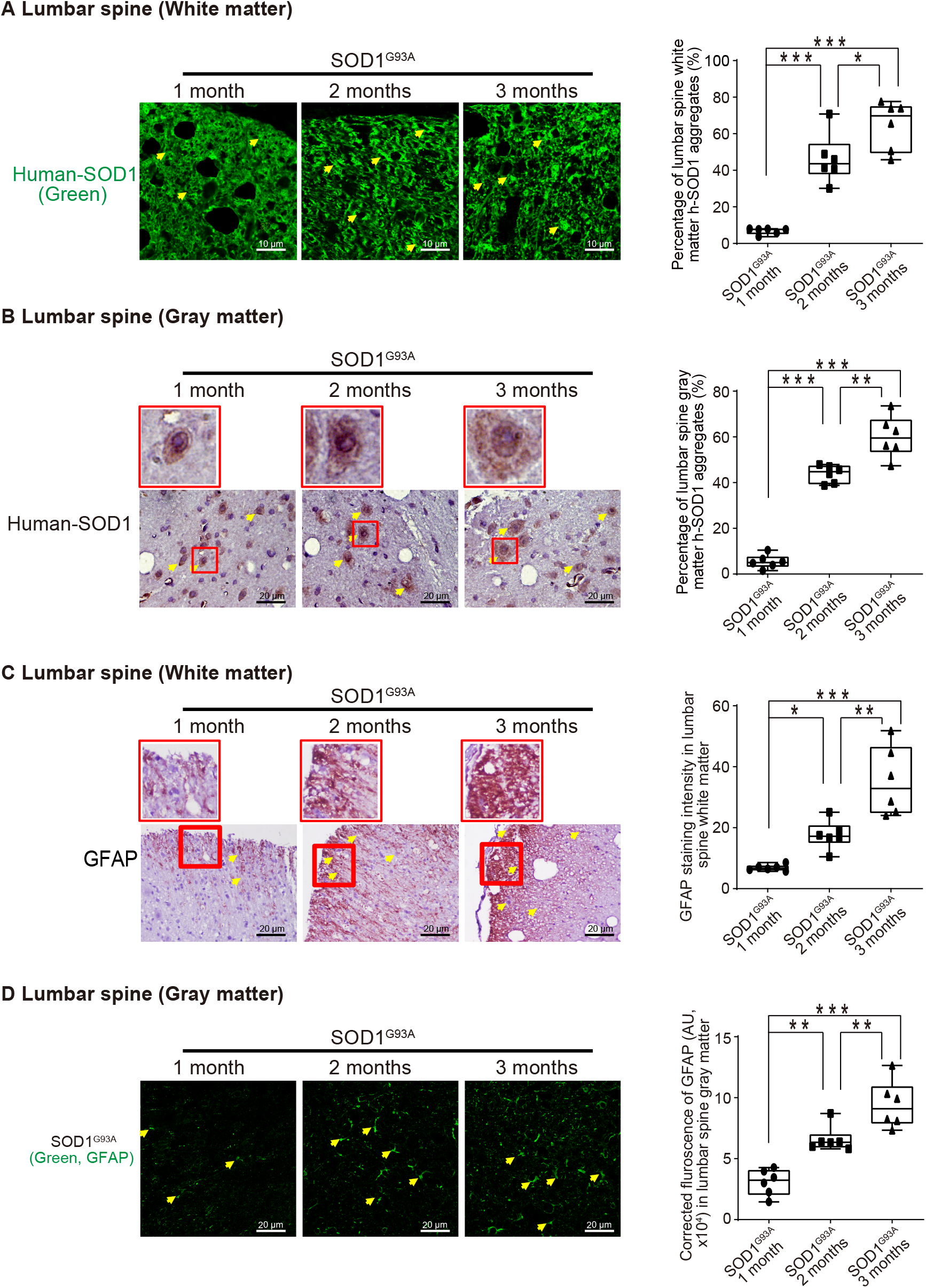
Human SOD1^G93A^ protein was aggregated, and GFAP protein was increased in the neurons during ALS progression. Aggregation of human-SOD1^G93A^ protein was observed in the lumbar spine white matter ***(A)*** and gray matter ***(B)*** starting in 2-month-old SOD1^G93A^ mice in immunofluorescence staining, immunohistochemistry staining and human-SOD1 protein aggregates percentage analysis. Images are from a single experiment, representative of 6 mice per group. (Data are expressed as mean ± SD. n = 6, one-way ANOVA test, *P < 0.05, **P < 0.01, ***P < 0.001). GFAP expression was enhanced starting from 2-month-old in the lumbar spine white matter ***(C)*** and gray matter ***(D)*** in the SOD1^G93A^ mice in immunofluorescence staining, immunohistochemistry staining and GFAP staining quantification. Images are from a single experiment, representative of 6 mice per group. (Data are expressed as mean ± SD. n = 6, one-way ANOVA test, *P < 0.05, **P < 0.01, ***P < 0.001).

### Association of altered ENS and the increased aggregation of SOD1G93A in G93A mice in longitudinal studies

Our longitudinal data at the functional, cellular, and protein levels showed the reduced intestinal motility, weak muscle strengths, altered ENS markers, and enhanced aggregation of SOD1^G93A^ starting at the 2-month-old (**Fig. S1**). Furthermore, we did correlation data analysis and showed that slow intestinal motility significantly correlated with weaker muscle strength, the changes of ENS marker GFAP, and an increased SOD1^G93A^ aggregation in the colon of the 2-month-old G93A mice (Fig. 4A). Immunohistochemistry staining and intestinal human-SOD1 ^G93A^ protein aggregates was further analyzed. We showed that a significant increase of human SOD1^G93A^ in the colon of the 2-month-old transgenic G93A mice were correlated with the decreased muscle strength (e.g., latency to fall and grip strength), decreased SMMHC, and increased GFAP, respectively (Fig. 4B). Decrease of SMMHC significantly linked with an increase of GFAP and PGP9.5 in the 2-month-old SOD1^G93A^ mice (Fig. 4C). Moreover, the changes of SMMHC and GFAP, from 3-month-old ALS colon, were correlated with slower intestinal movement, weaker muscle strength, and increased SOD1^G93A^ aggregation in the G93A mice (**Fig. S2**).

**Figure 4.**
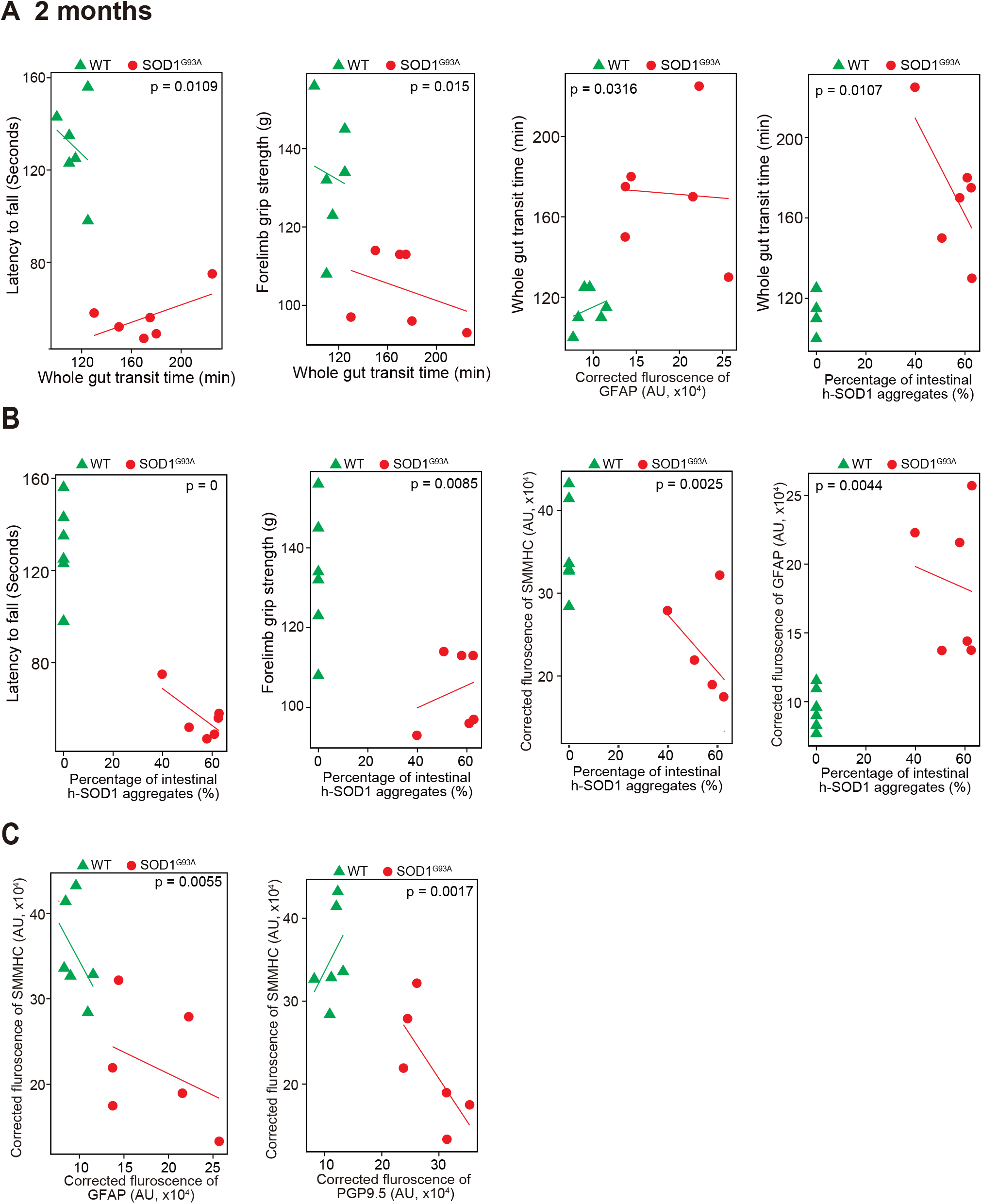
Association of altered ENS and increased aggregation of human-SOD1^G93A^ in the SOD1^G93A^ mice in longitudinal studies. ***(A)*** The correlation analysis between intestinal motility (Whole gut transit time) and latency to fall, forelimb grip strength; staining intensity of GFAP, aggregation of human SOD1^G93A^ protein respectively in the 2-month-old SOD1^G93A^, compared with WT mice. (P values are labeled in figures, n = 6). Slow Intestinal motility links with decreased forelimb grip strength, increased staining intensity of GFAP, increased aggregation of human SOD1^G93A^ protein respectively in 2-month-old SOD1^G93A^. ***(B)*** The correlation analysis between aggregation of human SOD1^G93A^ protein and latency to fall, forelimb grip strength, staining intensity of SMMHC, staining intensity of GFAP protein respectively in 2-month-old SOD1^G93A^, compared with WT mice. (P values are labeled in figures, n = 6). Increased aggregation of human SOD1^G93A^ protein links with decreased forelimb grip strength, decreased SMMHC, increased GFAP in the 2-month-old SOD1^G93A^. ***(C)*** The correlation analysis between staining intensity of SMMHC and staining intensity of GFAP, staining intensity of PGP9.5 protein respectively in the 2-month-old SOD1^G93A^, compared with WT mice. (P values are labeled in figures, n = 6). Decreased staining intensity of SMMHC links with increased staining intensity of GFAP and PGP9.5 respectively in the 2-month-old SOD1^G93A^, compared with WT mice.

### G93A mice with butyrate treatment showed a significantly longer latency to fall in the rotarod test, reduced SOD1^G93A^ aggregation, and decreased GFAP expression

Our previous study has shown the beneficial role of butyrate in delaying the progress of ALS ^8^. Here, we investigated the role of butyrate treatment on muscle strength and ENS. Male and female SOD1^G93A^ mice were treated with or without 2% butyrate in the drinking water starting from 63 days. SOD1^G93A^ mice with butyrate treatment showed significantly higher body weight than the mice without treatment (Fig. 5A). At age 14-15 weeks, the mice were subjected to the trial on the accelerating spindle (4 to 40 rpm) for 300 seconds. Latency to fall was recorded when the mouse fell from the rod. Each mouse was tested in 4 trials per day for 2 consecutive days. We found that G93A mice with butyrate treatment had a significantly longer latency to fall in the rotarod test, compared to the G93A mice without treatment (Fig. 5B). Butyrate treatment led to increased SMMHC and decreased GFAP expression in SOD1^G93A^ mice (Fig. 5C). We also found the reduced human-SOD1^G93A^ aggregation, decreased GFAP expression, and enhanced SMMHC expression in the intestines of SOD1^G93A^ mice with butyrate treatment (Fig. 5D). In the lumbar spine of SOD1^G93A^ mice with a butyrate treatment, we found reduced human-SOD1^G93A^ aggregation (Fig. 5E) and decreased GFAP expression (Fig. 5F). Taken together, these data suggested that butyrate treatment led to a significantly extended latency to fall in the rotarod test, reduced SOD1^G93A^ aggregation, and decreased GFAP expression.

**Figure 5.**
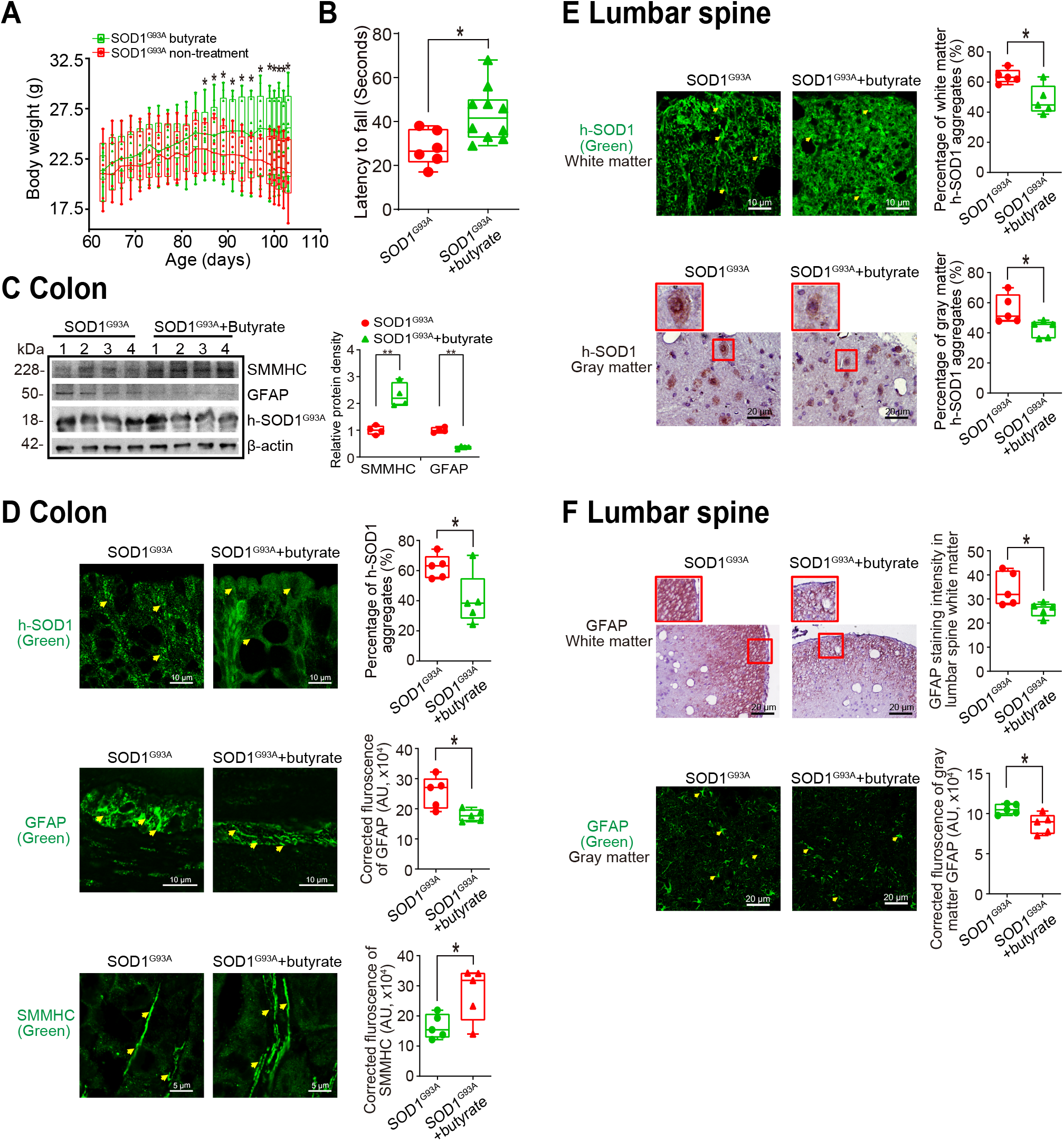
Butyrate treatment led to reduced human-SOD1^G93A^ aggregation, and enhanced ENS and muscle function in intestine and neurons of SOD1^G93A^ mice. ***(A)*** Body weight changes of SOD1^G93A^ mice after butyrate treatment. Male or female SOD1^G93A^ mice were treated with or without 2% butyrate in the drinking water starting from 63 days to 101 days. Butyrate treated SOD1^G93A^ mice started to show less weight loss from the age of 83 days, compared to the no-treatment SOD1^G93A^ mice. Each data point represents the average body weight. (Data are expressed as mean ± SD. n = 6-10, two-way ANOVA test, *P < 0.05). ***(B)*** SOD1^G93A^ mice with butyrate treatment showed a significantly increased rotarod test time. At age 14-15 weeks, the mice were subjected to the trial on the accelerating spindle (4 to 40 rpm) for 300 seconds. Latency to fall was recorded when the mouse fell from the rod. Each mouse was tested in 4 trials per day for 2 consecutive days. The mean times for 8 trials of the tests were calculated for each mouse. (Data are expressed as mean ± SD. n = 6-10, Welch’s t-test, *P < 0.05). ***(C)*** Butyrate treatment led to increased SMMHC and decreased GFAP expression in SOD1^G93A^ mice. (Data are expressed as mean ± SD. n = 4, student t-test, **P < 0.01). ***(D)*** Reduced human-SOD1^G93A^ aggregation, decreased GFAP expression and enhanced SMMHC expression in intestine of SOD1^G93A^ mice with butyrate treatment. Images are from a single experiment and are representative of 5 mice per group. (Data are expressed as mean ± SD. n = 5, Welch’s t-test, *P < 0.05). ***(E)*** Reduced human-SOD1^G93A^ aggregation in lumbar spine of SOD1^G93A^ mice with butyrate treatment. Images are from a single experiment and are representative of 5 mice per group. (Data are expressed as mean ± SD. n = 5, Welch’s t-test, *P < 0.05). ***(F)*** Decreased GFAP expression in lumbar spine of SOD1^G93A^ mice with butyrate treatment. Images are from a single experiment and are representative of 5 mice per group. (Data are expressed as mean ± SD. n = 5, Welch’s *t*-test, *P < 0.05).

### Antibiotics treatment led to reduced human-G93A SOD1 aggregation and enhanced ENS and muscle function in the intestine and neurons of SOD1^G93A^ mice

To test the role of manipulating microbiome in ALS, male and female G93A mice were also treated with antibiotics 33 (1 mg/ml metronidazole and 0.3 mg/ml clindamycin) in the drinking water starting from 48 day-old. The antibiotics treated mice had less weight loss, compared to mice without treatment (Fig. 6A). At age 107 days, the mice were subjected to the trial on the accelerating spindle. Latency to fall was significantly longer with antibiotic treatment (Fig. 6B). Furthermore, antibiotics treatment led to increased SMMHC and decreased GFAP expression in SOD1^G93A^ mice, compared to the mice without treatment (Fig. 6C). We also found reduced human-SOD1^G93A^ aggregation, decreased GFAP expression, and enhanced SMMHC expression in the intestines of SOD1^G93A^ mice treated with antibiotics (Fig. 6D). We then tested human-SOD1^G93A^ aggregation and GFAP expression in lumbar spine of SOD1^G93A^ mice treatment with or without antibiotics. Antibiotic treatment restored the SMMHC (Fig. 6E) and reduced GFAP (Fig. 6F). These data suggest that altered intestinal microbiome and function correlate with the skeletal muscle activity and motor neuron function in ALS.

**Figure 6.**
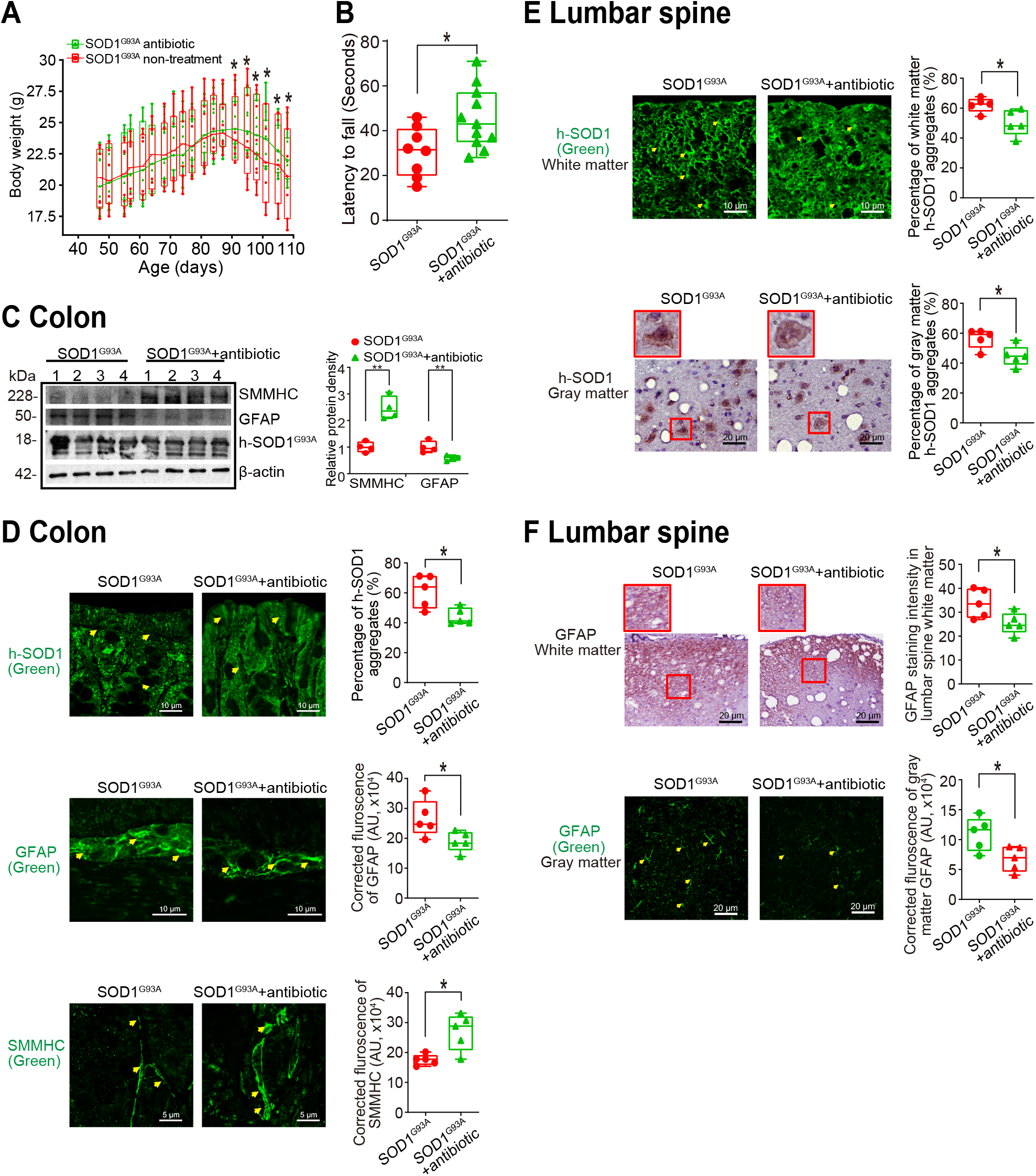
Antibiotics treatment led to reduced human-SOD1^G93A^ aggregation, and enhanced ENS and muscle function in intestine and neurons of SOD1^G93A^ mice. ***(A)*** Body weight changes of SOD1^G93A^ mice treated with antibiotics. Male or female SOD1^G93A^ mice were treated with or without antibiotics (1 mg/ml metronidazole and 0.3 mg/ml clindamycin) in the drinking water starting from 48 days to 107 days. Antibiotic treated SOD1^G93A^ mice started to show less weight loss at the age of 91 days, compared to the non-treatment SOD1^G93A^ mice. Each data point represents the average body weight. (Data are expressed as mean ± SEM. n = 8-11, two-way ANOVA test, *P < 0.05). ***(B)*** SOD1^G93A^ mice with antibiotics treatment showed a significantly increased time in the rotarod test. At 15 weeks old, the mice were subjected to the trial on the accelerating spindle (4 to 40 rpm) for 300 seconds. Latency to fall was recorded when the mouse fell from the rod. Each mouse was tested in 4 trials per day for 2 consecutive days. The mean times for 8 trials of the tests were calculated for each mouse. (Data are expressed as mean ± SD. n = 8-11, Welch’s t-test, *P < 0.05). ***(C)*** Antibiotics treatment led to increased SMMHC and decreased GFAP expression in SOD1^G93A^ mice. (Data are expressed as mean ± SD. n = 4, student t-test, *P < 0.05, **P < 0.01). ***(D)*** Reduced human-SOD1^G93A^ aggregation, decreased GFAP expression and enhanced SMMHC expression in intestine of SOD1^G93A^ mice treatment with antibiotic. Images are from a single experiment and are representative of 5 mice per group. (Data are expressed as mean ± SD. n = 5, Welch’s t-test, *P < 0.05). ***(E)*** Reduced human-SOD1^G93A^ aggregation in lumbar spine of SOD1^G93A^ mice treatment with antibiotics. Images are from a single experiment and are representative of 5 mice per group. (Data are expressed as mean ± SD. n = 5, Welch’s t-test, *P < 0.05). ***(F)*** Decreased GFAP expression in lumbar spine of SOD1^G93A^ mice treatment with antibiotics. Images are from a single experiment and are representative of 5 mice per group. (Data are expressed as mean ± SD. n = 5, Welch’s *t*-test, *P < 0.05).

### Increased aggregation of SOD1 mutant protein by fecal microbes in human colonoids

Colonoids derived from intestinal stem cells were used to study the host-microbial interactions, as we did in the previous studies ^34–37^. First, we transfected human colonoids with SOD1^G93A^-GFP plasmids after 48 hours, then treated with feces from WT or SOD1^G93A^ mice at different ages for 2 hours (Fig. 7). Colonoids were imaged overtime (Fig. 7A), then collected for immunostaining and western blots. We did not see significant changes of buds or size of organoids after treatment, as shown in Fig. 7A. Interestingly, we found that feces from the ALS mice were able to induce SOD1 aggregated in human colonoids transfected with SOD1^G93A^-GFP plasmids (Fig. 7B). More SOD1^G93A^ protein aggregations were induced by feces of 2- and 3-month-old SOD1^G93A^ mice (Fig. 7C), suggesting the direct effect of microbes in feces of SOD1^G93A^ mice to increase the protein SOD1^G93A^ aggregation. It also indicated the age-dependent functional differences of microbiome in the SOD1^G93A^ mice.

**Figure 7.**
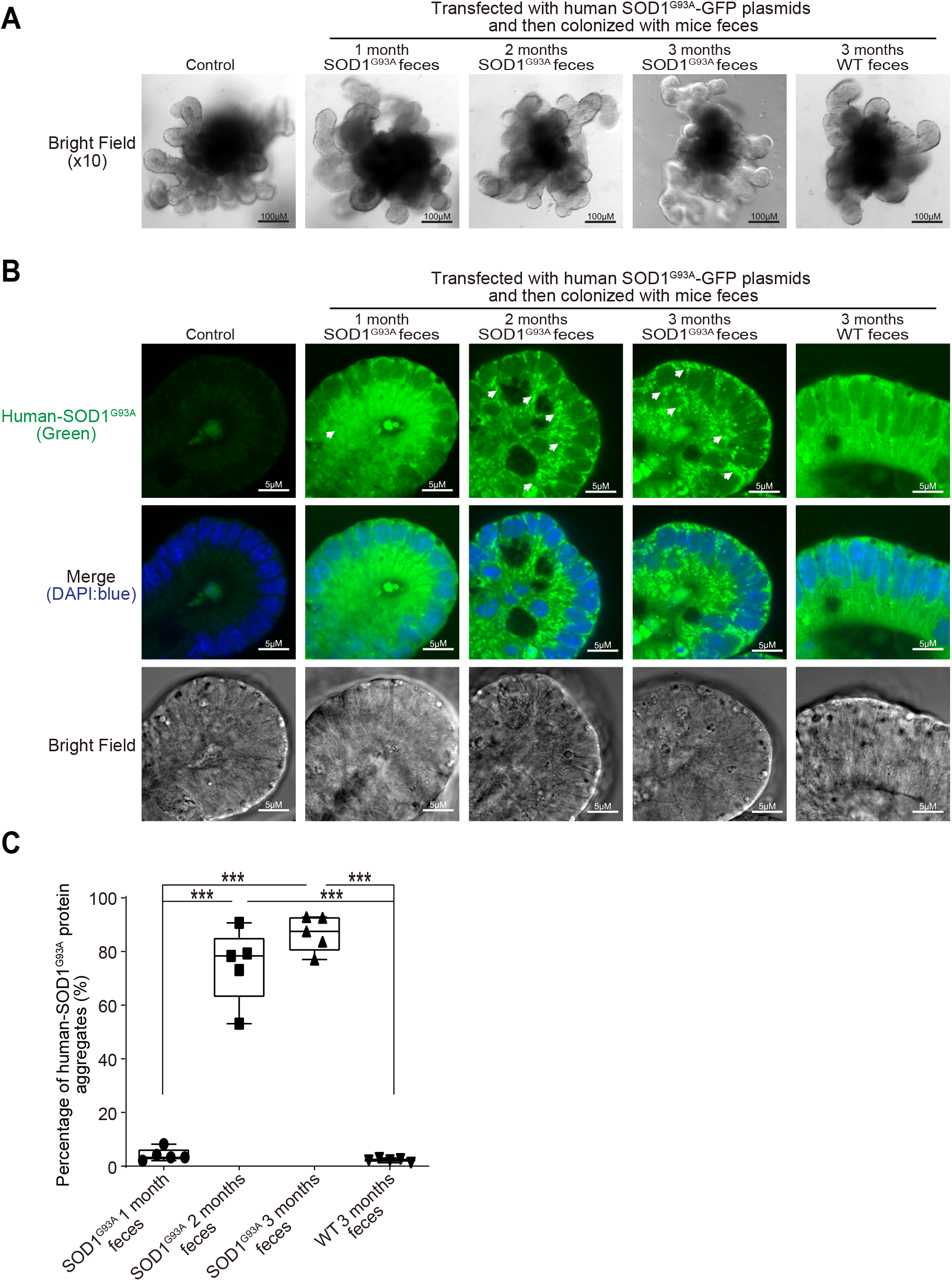
Increased aggregation of SOD1 mutant protein in human colonoids transfected with SOD1^G93A^-GFP plasmids and then colonized with SOD1^G93A^ or WT mice feces. **(*A*)** Human colonoids were transfected with SOD1^G93A^-GFP plasmids after 48 hours and treated with feces from SOD1^G93A^ / WT mice at different ages for 2 hours. After 2-hour treatment of mice feces, the phenotype of human colonoids, e.g., shape and buds did not change. Images are representative of experiments in triplicate. **(*B*)** Human SOD1 expression in human colonoids were transfected with SOD1^G93A^-GFP plasmids and then colonized with feces from SOD1^G93A^ or WT mice. Images are representative of experiments in triplicate. ***(C)*** More SOD1^G93A^ protein aggregations in the human colonoids after SOD1^G93A^-GFP plasmid transfection and 2 and 3-month-old SOD1^G93A^ mice feces colonization. (Data are expressed as mean ± SD. n = 5, one-way ANOVA test, ***P < 0.001).

### Changes of fecal microbiome in SOD1^G93A^ mice

To identify the changes of bacterial community in the feces of the SOD1^G93A^ mice, we performed the longitudinal data analysis of the microbiome data from 1-, 2-, and 3-month-old mice, compared with the age-matched WT mice. At the species level, we found the significantly different changes of certain bacteria between groups in a time-dependent **manner (**Table 1). We used different models (e.g., Fast Zero-inflated Negative Binomial Mixed Modeling^38^ and linear mixed-effects models ^39, 40^) to examine the changes of microbiome at the species level, as indicated in Table 1.

**Table 1:**
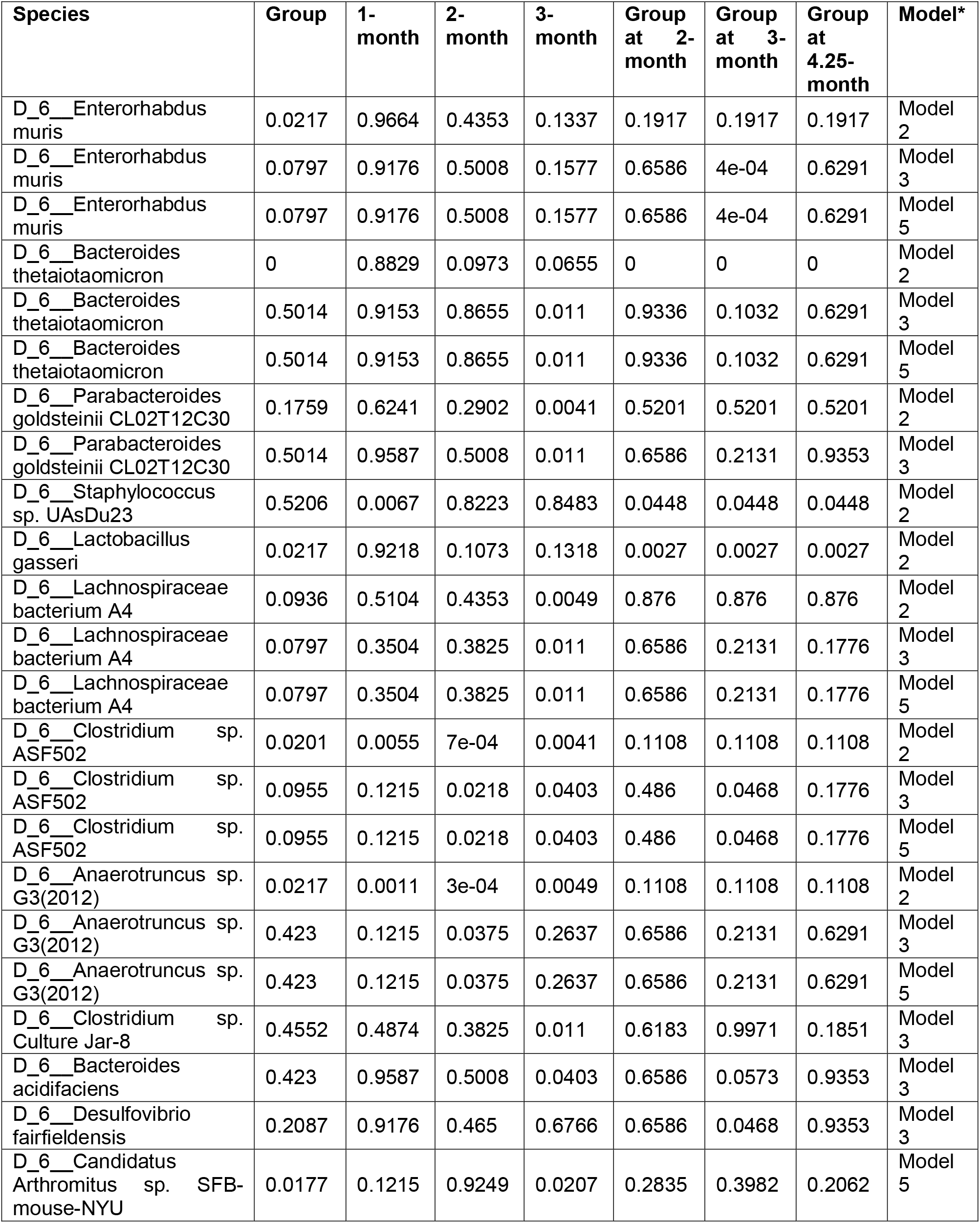
Longitudinal study for microbiome of SOD1^G93A^ mice and age-matched WT mice, comparing group, time and group and time interaction.

To confirm the identified significant changes of bacteria by the longitudinal models and visualize the changes over time, we performed the line plots (Fig. 8) and conducted ANOVA, Kruskal-Wallis rank sum test, Welch’s t-test or Wilcoxon test as appropriate for specific bacteria. Specifically, *Enterohabdus Muris* started increasing in the 2-month-old SOD^G93A^, compared with the 1-month-old SOD^G93A^ mice, and further significantly enhanced in the 3-month-old SOD1^G93A^ mice (Fig. 8A). *Staphylococcus. Sp. UAsDu23* was enhanced in 1-month-old SOD^G93A^ mice, and then significantly enhanced over time, compared with the age matched WT mice (Fig. 8B). *Clostridium sp. ASF502* was significantly reduced in the 3-month-old G93A, compared with WT mice (Fig. 8C). *Desulfovibrio fairfieldensis* started to increase in the 2-month-old SOD^G93A^, compared with 1-month old SOD^G93A^ mice and was significantly enhanced in the 3-month-old SOD1^G93A^ mice, compared with WT mice. In contrast, the relative abundance of *Desulfovibrio fairfieldensis* was very stable in the WT mice (Fig. 8D). The dramatic changes of bacteria in the SOD^G93A^ mice may explain the ability of the feces to enhance SOD1 aggregation in the colonoids.

**Figure 8.**
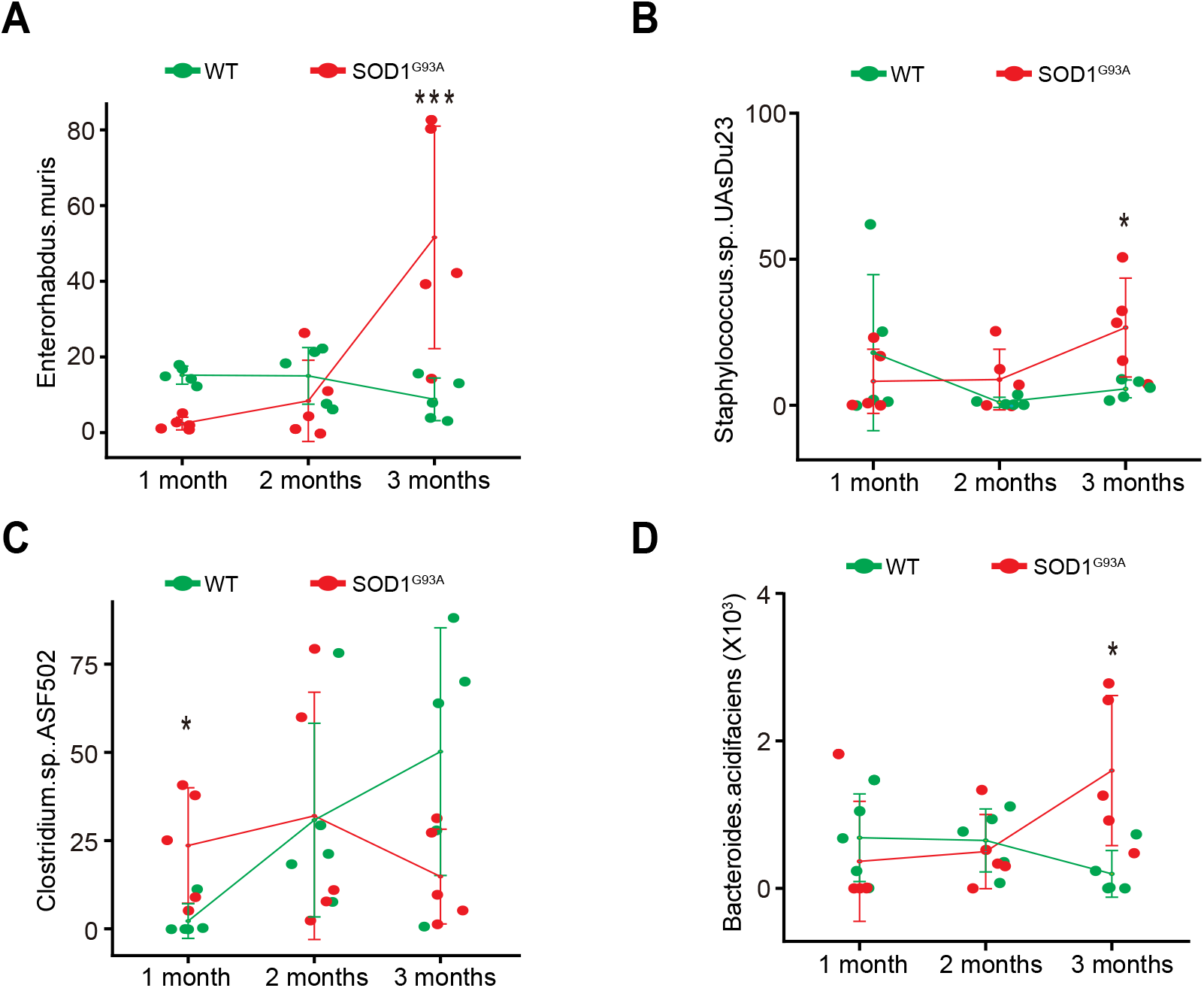
Fecal microbiome changes in SOD1^G93A^ mice. ***(A)*** *Enterohabdus Muris* started increasing in 2-month-old SOD^G93A^, and then significantly enhanced in 3-month-old SOD1^G93A^ mice, compared with WT mice. (n = 5, two-way Kruskal-Wallis rank sum test, ***P < 0.05 adjusted by FDR.). ***(B)*** *Staphylococcus. Sp. UAsDu23* started increase in 2-month-old SOD^G93A^, compared with 1-month-old SOD^G93A^ mice, and then significantly enhanced in 3-month-old SOD1^G93A^ mice, compared with WT mice. (n = 5, Welch’s t-test, *P < 0.05). ***(C)*** Clostridium sp. ASF502 enhanced in 1months SOD^G93A^, compared with WT mice (n = 5, Welch’s t-test, *P < 0.05). ***(D)*** *Desulfovibrio fairfieldensis* significantly enhanced in 3-month-old SOD1^G93A^ mice, compared with the WT mice. (n = 5, two-way ANOVA test, *P < 0.05, adjusted by the Tukey method).

### Butyrate treatment reduced the microbial differentiation between WT and G93A mice

Interestingly, the changes of fecal microbiome in SOD1^G93A^ mice were reduced after butyrate treatment (Table 2). For example, the significantly enhanced *Enterohabdus Muris* in 3-month-old SOD1G93A mice was not significant anymore after butyrate treatment, compared with WT mice (n = 6, two-way Kruskal-Wallis rank sum test, ***P < 0.2 adjusted by FDR.) The significantly enhanced *Staphylococcus. Sp. UAsDu23* in 3-month-old SOD1^G93A^ mice without butyrate treatment was not significant after butyrate treatment (n = 6, Welch’s t-test, P = 0.55). The identified Clostridium sp. ASF502 in 1-month-old SOD^G93A^, compared with 1-month-old WT mice, was no longer significant after butyrate treatment (n = 6, Wilcoxon rank sum test, P = 0.6). The significantly enhanced *Desulfovibrio fairfieldensis* in the 3-month-old SOD1^G93A^ mice was no longer significant, compared with the WT mice (n = 6, Kruskal-Wallis rank sum test, P = 0.4 adjusted by the FDR method.). Instead, *Lachnospiraceae bacterium* A4, which was not different without butyrate treatment in the 3-month-old SOD1^G93A^ mice, was significantly enhanced after butyrate treatment. Without butyrate, at 1-month-old, *Candidatus Arthromitus sp. SFB.mouse.NYU*, a bacterium in regulating Th17 immunity ^41^, in the SOD1^G93A^ mice was significantly different from the WT, but with butyrate treatment, is not significant difference between. These data indicate the beneficial roles of butyrate may restore the microbiome and reduce the significant difference between WT and SOD1^G93A^ mice, thus delaying the ALS progress of SOD1^G93A^ mice.

**Table 2:** Longitudinal study for microbiome of SOD1^G93A^ mice and age-matched WT mice with butyrate treatment, comparing group, time and group and time interaction.

| Species | Group | 1-month | 3-month | Group at 1-month | Group at 3-month | Model* |
| --- | --- | --- | --- | --- | --- | --- |
| D_6__Parabacteroides goldsteinii CL02T12C30 | 0.777 | 0.002 | 0.609<br>3 | 0.1302 | 0.867 | Model 1 |
| D_6__Parabacteroides goldsteinii CL02T12C30 | 0.9366 | 0.0106 | 0.928 | 0.1848 | 0.9547 | Model 2 |
| D_6__Clostridium sp. ASF356 | 0.6512 | 0.287 | 0.016<br>4 | 0.7647 | 0.1022 | Model 1 |
| D_6__Clostridium sp. ASF356 | 0.5862 | 0.2137 | 0.024 | 0.7731 | 0.1868 | Model 2 |
| D_6__Clostridium sp. ASF502 | 0.7548 | 0.003 | 0.001<br>1 | 0.7647 | 0.867 | Model 1 |
| D_6__Clostridium sp. ASF502 | 0.9366 | 0.0713 | 0.020<br>4 | 0.8169 | 0.7351 | Model 2 |
| D_6__Clostridium sp. Culture Jar-8 | 0.0826 | 0.219 | 0.806<br>2 | 0.056 | 0.3042 | Model 1 |
| D_6__bacterium enrichment culture clone M244 | 0.9366 | 0.3992 | 0.044<br>3 | 0.9329 | 0.7351 | Model 2 |

\* Model 1 was fitted by Fast Zero-inflated Negative Binomial Mixed Modeling (FZINBMM) <sup>38</sup> using group, time and their interaction term as fixed effects and intercept as random effect. The within-subject correlation of AR(1) was used to account for within-individual correlation and the log(total read counts in sample) was used as the offset to adjust for library sizes.
Model 2 was fitted by the linear mixed-effects models <sup>39, 40</sup> using group, time and their interaction term as fixed effects and intercept as random effect. The within-subject correlation of AR(1) was used to account for within-individual correlation and the arcsine square root transformation of taxa abundance counts was used in lieu of normalization.
Model 3 was fitted by the linear mixed-effects models <sup>39, 40</sup> using group, time and their interaction term as fixed effects and intercept as random effect. The arcsine square root transformation of taxa abundance counts was used in lieu of normalization, but the independence of correlation matrix was used to assume the most simplified within-individual structure.

## Discussion

In the current study, we have demonstrated a novel link between intestinal mobility, ENS, and microbiome in the SOD1 aggregation and progression of ALS. Dysbiosis occurred at the early stage of the G93A mice (1-month-old) before observed dysfunction of ENS. The timeline for the changes of microbiome, ENS, SOD1 aggregation in different organs, and muscle strength indicate the microbial difference in the 1-month-old ALS mice, compared to the WT mice (Fig. 9). The G93A mice aged one-month-old showed no significant changes of ENS and motility but altered gut microbiome (e.g. increased *Clostridium sp. ASF502*), compared with the WT mice. At 2-month-old before ALS onset, G93A mice had significant lower intestinal motility, decreased grip strength, and reduced time in the rotarod. These changes correlated with consistent dysbiosis and increased aggregation of mutated SOD1^G93A^ in the colon and small intestine. We observed increased GFAP and decreased SMMHC expression. At the ALS onset period (3 months), G93A mice had much slower intestinal motility, decreased grip strength, reduced time in rotarod test. GFAP expression was increased and SMMHC decreased. Human-G93A-SOD1 mutated protein showed severe aggregation in the intestine. Moreover, we found that butyrate treatment and antibiotic treatment restored some of the intestinal mucosal functions and corrected dysbiosis in the ALS mice. Manipulating the microbiome improves the muscle performance of G93A mice. Our study provides insights into fundamentals of intestinal neuromuscular structure/function and microbiome in ALS.

**Figure 9.**
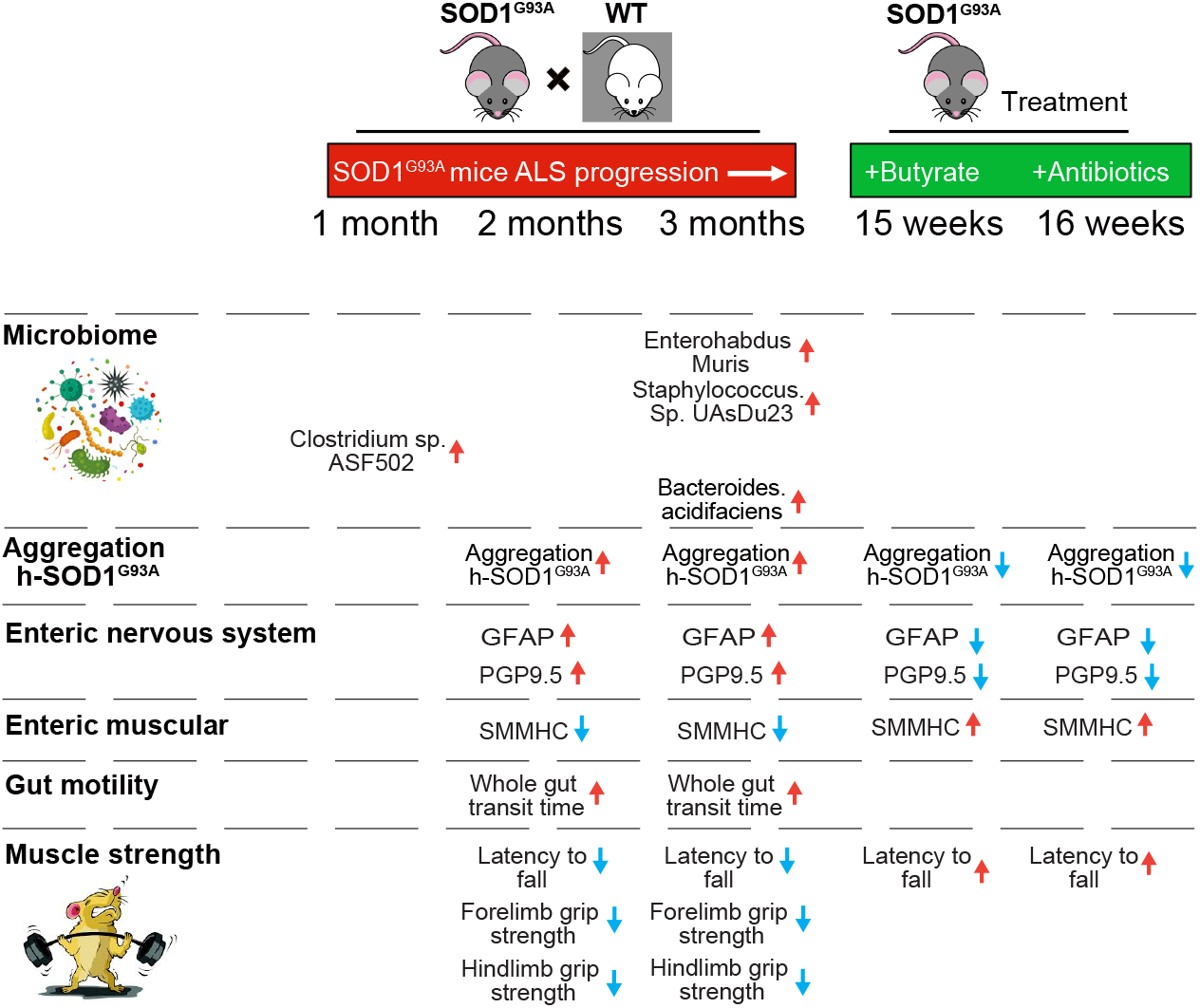
Timeline of altered microbiome, SOD1^G93A^ aggregation, EN function, and lumbar spine pathophysiological in the progression of SOD1^G93A^ ALS mice.

Our longitudinal studies in the ALS human-SOD1^G93A^ mice have revealed microbial communities closely tied to host metabolism and immunity in ALS. We have previously reported the significant reduction of butyrate-producing bacteria, e.g., Roseburia (family *Lachnospiraceae*, order *Clostridiales*), in the SOD1^G93A^ mice ^8, 9^. Butyrate-producing bacteria are known to play an important role in the control of gut inflammatory processes and in the maturation of the immune system, primarily through the production of butyrate ^42^. Here, at the species level, we further identified bacteria that are important for the immunity and inflammatory response. For example, *Enterohabdus Murisi* was first isolated from an IBD model ^63^*. Lachnospiraceae bacterium A4* bacteria is reported to inhibit Th2-cell differentiation by inducing dendritic cell production of TGF-β.^43^ *Clostridium sp. ASF502. Clostridium sp. ASF502* belongs to altered Schaedler flora ^44^(ASF), a model microbial community with both in vivo and in vitro relevance and are known important for the ASF mice to fully develop immune system, resistance to opportunistic pathogens, and normal GI function and health. *Candidatus Arthromitus* sp. SFB-mouse-NYU is among the segmented filamentous bacteria (SFB) direct the accumulation of potentially proinflammatory Th17 cells in the intestinal lamina propria ^41^. *Candidatus Arthromitus.sp. SFB.mouse.NYU* in the 1-montn-old G93A mice without butyrate, is significant different from the WT mice, but with butyrate treatment, there was not significant any more between the G93A and WT mice. *Desulfovibrio fairfieldensis* is also reduced after the butyrate treated G93A mice. *Desulfovibrio fairfieldensis* is among the sulfate-reducing bacteria, which are anaerobic microorganisms that conduct dissimilatory sulfate reduction to obtain energy, resulting in the release of a great quantity of sulfide. Altered sulfate-reducing bacteria are reported in human IBD cases ^64^. Taken together, our data suggested the early ALS-associated dysbiosis of intestinal microbiome related to host metabolism and immunity and microbial manipulation by butyrate or antibiotics could slow down the disease progression.

Here, we define the changes in intestinal microbiome and function during ALS progression, and their correlation with the progressive decline of skeletal muscle activity and motor neuron. A previous study has shown correlations between bacterial species abundances and transit times are diet dependent^18^. To understand how different diets, microbiota, and the ENS interact to regulate gut motility, this study used a gnotobiotic mouse model that mimics short-term dietary changes that happen when humans are traveling to places with different culinary traditions. The levels of unconjugated bile acids-generated by bacterial bile salt hydrolases-correlated with faster transit, including during consumption of a Bangladeshi diet. This study demonstrated how a single food ingredient interacts with a functional microbiota trait to regulate intestinal motility ^18^. Here, we showed that butyrate, a bacterial product, significantly restored the ENS and smooth muscle in the ALS mice. Manipulating the microbiome by antibiotics also facilitates the ENS and muscle, thus delaying ALS onset and progression. An ALS study also indicates the temporal evolution of the microbiome, immune system, and epigenome with the disease progression of the SOD1^G93A^ mice ^48^. Our data in the ALS model and human colonoids have provided novel links among microbiome, SOD1 aggregation, and intestinal dysfunction. Dysbiosis and SOD1 mutated aggregation occurred at the early stage of the G93A mice before observed slow intestinal motility and dysfunction of ENS, suggesting a biomarker for ALS diagnosis. Our data further suggest that restoring a healthy microbiome reduced aggregation of the SOD1 G93A-mutated protein in the intestine and nervous tissues and slowed down the progression of ALS in mice.

On one hand, it is unknown whether intestinal epithelial damage directly affects the enteric neurons and smooth muscles. On the other hand, the ENS is an important regulator of the proliferation and differentiation of the mucosal epithelium ^47^. Interestingly, the transgenic mice with human ALS mutation Prp-TDP43 A315T ^46^ had a defect in enteric neurons, which may be associated with microbiome and mucosal barrier abnormalities. In the future study, we will investigate the role of other ALS risk genes on the microbiome and ENS. The mechanism of motility dysfunction depends on the intestinal region and the stage of diseases ^49, 50^. There is an age-dependent shift in macrophage polarization that causes inflammation-mediated degeneration of ENS ^51^. Further studies on pre-inflammatory serum cytokines and LPS in the patients combined with the changes of microbiome will help with the early diagnosis of the disease.

Sporadic (SALS) and familial (FALS) forms of ALS manifest similar pathological and clinical phenotypes, suggesting that different initiating causes lead to a mechanistically similar neurodegenerative pathway. Some clinical studies have indicated intestinal abnormalities in ALS patients ^52–54^. Increased LPS is reported in SALS patients ^5, 55^. ALS patients have elevated intestinal inflammation and dysbiosis ^10, 11^. The therapeutic methods to target microbiome and intestinal functions will help both SALS and FALS patients.

Currently, treatment with the FDA approved drug, Riluzole, merely extends the patient’s life span for a few months. The second new drug Radicava was approved in 2017. However, there is still a need to develop new treatments for alleviating disease progression and improving the life quality of ALS patients. A phase 2 randomized, placebo-controlled trial treated patients for 24 weeks, which involved 137 ALS patients, 89 of whom were treated with the combination of sodium phenylbutyrate plus taurursodiol ^56^. It found a modest reduction in functional decline in patients receiving the combination therapy ^56^, suggesting the promising role of sodium phenylbutyrate. However, this study has no data on the changes of the human microbiome. A recent study reported the microbiota changes as a potential human ALS biomarker and indicated the microbial strategy could be considered to restore the status of the intestine ^14^. A larger trial testing more patients over a longer period is also needed.

The limit of the current study is that there is no ENS data in human ALS. The direct interactions between ENS and microbiome are unknown. Although human ALS and ENS functions have yet to be investigated, the insights gained from the analysis of the ALS mice and microbiome will contribute to a better understanding of how microbiome affect the host in the progression of a disease. The direct role of other microbial metabolites, in addition to butyrate, on the function of ENS and its mechanisms are also in the future plan.

## Conclusion

We have demonstrated a novel link between microbiome, SOD1 aggregation, and intestinal mobility. Dysbiosis at the early stage of the G93A mice occurs before observed dysfunction of enteric neuromuscular system. SOD1 mutated aggregation occurred in both intestine and spinal cord. Interestingly, fecal samples from the G93A mice were able to induce SOD1 mutated aggregation in human colonoids. Targeting the intestinal microbiome was able to improve neuromuscular function in ALS. Our study provides insights into fundamentals of intestinal neuromuscular structure/function and microbiome in ALS, suggesting the potential to use microbial biomarkers for the diagnosis and to manipulate the intestinal microbiome for the treatment.

## Materials and Methods

### Animals

SOD1^G93A^ and age matched wild type mice were used in this study, SOD^G93A^ mice were originally purchased from Jackson Laboratory (B6SJL-Tg (SOD1-G93A) 1Gur/J, stock No.002726). All experiments were carried out in strict accordance with the recommendation in the Guide for the Care and Use of Laboratory Animals of the National Institutes of Health. Mice were provided with water ad libitum and maintained on a 12-hour dark/light cycle. The protocol was approved by the IACUC of University of Illinois at Chicago Committee on Animal Resources (ACC 18-233).

### Measurement of gastrointestinal transit times

On the test day, mice were transferred to individual empty plastic cages (devoid of bedding) and were deprived of food, with free access to water. Two hours after food deprivation, mice were then gavaged with 150 μl of Evans blue marker (5% Evans blue, 5% gum Arabic in drinking water) between 09:00 and 09:30 local time. After gavage, mice were fed ad libitum. The mice were observed at 5 min intervals until feces with blue were eliminated (maximum time of observation was 450 min). The time from the end of gavaging to the first blue faecal pellet was measured in minutes and constituted the whole gut transit time.

### Rotarod test

Motor coordination, strength and balance were assessed using a rotarod (Harvard Apparatus, Holliston, MA). Mice started to train on the rotarod daily three days before recording the data. Animals were placed onto the rod at a speed of 4 rpm which accelerates over the course of 300 seconds to 40 rpm. Latency to fall was recorded when the mouse fell from the rod. Each mouse was tested in 4 trials per day for 2 consecutive days. The mean times for 8 trials of the tests were calculated for each mouse.

### Assessment of grip strength

Forelimb and hindlimb grip measurements were acquired in triplicate with a 25 N Grip strength meter (Harvard Apparatus, Holliston, MA, USA). The mice were lowered onto a triangle bar of the grip strength meter until the animals gripped the bar with their forelimbs or hindlimbs, then the mice were pulled gently backward until they released their grip. The force gauge of the grip meter recorded the maximum force.

### Western blot analysis and antibodies

Mouse intestinal mucosal cells were collected by scraping from mouse colon, including proximal and distal colon, as previously described. ^57^ Briefly, mouse mucosal cells were lysed in lysis buffer (1% Triton X-100 (Sigma-Aldrich, X100), 150 mM NaCl (J.T. Baker 3624-19), 10 mM Tris (Fisher Scientific, BP152-5) pH 7.4, 1 mM EDTA (Fisher Scientific, BP120-1), 1 mM EGTA (Sigma-Aldrich, E3889) pH 8.0, 0.2 mM sodium ortho-vanadate (Sigma-Aldrich, S6508) and protease inhibitor cocktail (Roche Diagnostics, 118367001). Cultured cells were rinsed twice in ice-cold Hanks’ balanced salt solution (Sigma-Aldrich, H1387), lysed in protein loading buffer (50 mM Tris, pH 6.8, 100 mM dithiothreitol (Amresco, 0281), 2% SDS (Sigma-Aldrich, L3771), 0.1% bromophenol blue (IBI Scientific, IB74040) and 10% glycerol (Sigma-Aldrich, G5516)) and sonicated (Branson Sonifier, 250). Equal amount of protein was separated by SDS-polyacrylamide gel electrophoresis, transferred to nitrocellulose (Bio-rad, 162-0112) and immunoblotted with primary antibodies: anti-GFAP (Abcam, ab53554), SMMHC (Abcam, ab53219), human SOD-1^58^ or β-actin (Sigma-Aldrich, A1978) antibodies and visualized by ECL chemiluminescence (Thermo Scientific, 32106). Membranes probed with more than one antibody were stripped before re-probing. Western blot bands were quantified using Image Lab 4.01 (Bio-Rad).

### Immunofluorescence

Intestinal and Lumbar spine tissues were freshly isolated and paraffin-embedded after fixation with 10% neutral buffered formalin. Immunofluorescence was performed on paraffin-embedded sections (5 μm). After preparation of the slides as described previously^57^, tissue samples were incubated with anti-GFAP (Abcam, ab53554), SMMHC (Abcam, ab53219), α-SMA (Abcam, ab5694), PGP9.5 (Invitrogen, MA1-90008), human SOD-1 at 4°C overnight. Samples were then incubated with sheep anti-goat Alexa Fluor 594 (Life Technologies, A11058), or goat anti-mouse Alexa Flour 488 (Life Technologies, A-11001) and DAPI (Life Technologies, D1306) for 1 h at room temperature. Tissues were mounted with SlowFade (Life Technologies, s2828), followed by a coverslip, and the edges were sealed to prevent drying. Specimens were examined with Zeiss laser scanning microscope (LSM) 710. Fluorescence intensity was determined by using Image J software. This method determines the corrected total fluorescence by subtracting out background signal, which is useful for comparing the fluorescence intensity between cells or regions.

### Immunohistochemistry

After preparation of the slides, antigen retrieval was achieved by incubation of the slides for 15 min in the hot preheating sodium citrate (pH 6.0) buffer, and 30 min of cooling at room temperature. Endogenous peroxidases were quenched by incubating the slides in 3% hydrogen peroxide for 10 min, followed by three rinses with HBSS, and incubation for 1 hour in 3% BSA + 1% goat serum in HBSS to reduce nonspecific background. Primary antibodies anti-GFAP (Abcam, ab53554) or human SOD-1 were applied overnight in a cold room. After three rinses of the slides with HBSS, they were incubated in secondary antibodies (1:100, Jackson ImmunoResearch Laboratories, 115-065-174) for 1 hour at room temperature. After washing with HBSS for 10 minutes, the slides were incubated with vectastain ABC reagent (Vector Laboratories, PK-6100) for 1 hour. After washing with HBSS for five minutes, color development was achieved by applying a peroxidase substrate kit (Vector Laboratories, SK-4800) for 2 to 5 minutes, depending on the primary antibody. The duration of peroxidase substrate incubation was determined through pilot experiments and was then held consistently for all of the slides. After washing in distilled water, the sections were counterstained with haematoxylin (Leica, 3801570), dehydrated through ethanol and xylene, and cover slipped using a permount (Fisher Scientific, SP15-100). Immunohistochemistry intensity was also determined by using Image J software.

### Butyrate treatment in mice

SOD1^G93A^ and age matched wild type mice were divided into groups randomly. The butyrate-treated group receives sodium butyrate (Sigma-Aldrich, 303410) at a 2% concentration in filtered drinking water. Control group receives filtered drinking water without sodium butyrate. All animals are weighted and received detailed clinical examination, which included appearance, movement and behavior patterns, skin and hair conditions, eyes and mucous membranes, respiration and excreta. If restricted outstretching of the hind legs is observed when the tail is held, it means the symptom of ALS is obvious. Lie the mouse on the back, if it can’t turn over in 30 seconds, the mouse is humanely sacrificed.

### Antibiotic treatment in mice

SOD1^G93A^ and age matched wild type mice were divided into groups randomly. The antibiotic-treated group receives antibiotics (1 mg/ml metronidazole and 0.3 mg/ml clindamycin) in filtered drinking water. Control group receives filtered drinking water without antibiotics. All animals are weighted and received detailed clinical examination, which included appearance, movement and behavior patterns, skin and hair conditions, eyes and mucous membranes, respiration and excreta. If restricted outstretching of the hind legs is observed when the tail is held, it means the symptom of ALS is obvious. Lie the mouse on the back, if it can’t turn over in 30 seconds, the mouse is humanely sacrificed.

### Human colonoid transfection and treatment With SOD1^G93A^ mice feces

Human organoids were developed using endoscopy samples in the UIC hospital.

Crypts were released from colon tissue by incubation for 30□minutes at 4°C in phosphate-buffered saline containing 2□mmol/L EDTA. Isolated crypts were counted and pelleted. A total of 500 crypts were mixed with 50 L Matrigel (BD Biosciences, San Jose, CA) and plated in 24-well plates. The colonoids were maintained in Human IntestiCult Organoid Growth Medium (STEMCELL Technologies, Inc, Vancouver, BC). The human colonoids were transfected with the plasmids for fluorescent protein tagged human ALS-causing mutation SOD1^G93A^-GFP. Forty-eight hours after transfection, the human organoids were treated with SOD1^G93A^ mice feces.

Fresh feces were collected from 5 SOD1^G93A^ mice and then well mixed. A total of 100 mg feces was homogenized in 6 mL Hanks Balanced Salt Solution and centrifuged for 30 seconds at 300 rpm, at 4°C, to pellet the particulate matter. Organoids were treated with 250 μL feces supernatant for 2 hours, the organoids were washed 3 times with Hanks Balanced Salt Solution, and then the cells were incubated in regular organoid culture medium for 2 hours. The percentage of cells with human-SOD1 protein aggregates was counted in each experimental group.

### Microbiome 16S rRNA sequencing and Bioinformatic analysis

#### Study design for microbiome data

We designed two randomized controlled longitudinal microbiome studies for this project: without butyrate treatment and with butyrate treatment. The study without butyrate treatment was designed to collect fecal samples of G93A and wild type mice over mouse ages of 1-, 2-, 3- and 4.25-month-old. It aimed to access the genetic effects on the dynamics of gut microbes. The study with butyrate treatment was designed to collect fecal samples of G93A and wild type mice over mouse ages of 1- (pre-treatment), 2- and 3-month-old. It aimed to access the butyrate treatment effects on the dynamics of gut microbes.

#### Fecal sample collection, sampling

We collected fecal samples for 16S rRNA sequencing. Fresh fecal samples from each group (G93A and WT mice: for each time point, n = 5 from male and female mice) were collected from the colon and placed into the sterile tubes. The samples were kept with dry ice for low temperature and were sent to the University of Illinois at Chicago Research Resources Center for genomic sequencing. The genomic DNAs of samples were extracted using DNeasy Power Fecal Kit (12830, Qiagen, Hilden, Germany) based on manufacturer’s instructions with a slight modification.

The samples were heated at 65°C for 10 min before homogenizing with FastPre-24 5G bead-beating device (MP Biomedicals, Solon, OH, USA) at 6 m/s for 40 s. Genomic DNAs were fragmented into relatively small pieces (generally 250-600 bp fragments) before sequencing.

#### 16S rRNA sequencing

Genomic DNAs were PCR-amplified with the Earth Microbiome Project primers CS1_515F and CS2_806R targeting the V3–V4 regions of the 16S rRNA gene. A two-stage “targeted amplicon sequencing” protocol as described in ^65^ was used to generate amplicons. Two independent PCR steps were conducted in the workflow for preparing samples for next-generation amplicon sequencing. In the first stage, PCR amplification was performed using primers containing CS1 and CS2 linkers (CS1_341F: 50-ACACTGACGACATGGTTCTACAGTGCCAGCMGCCGCGGTAA-30;

CS2_806R: 50-TACGGTAGCAGAGACTTGGTCTGGACTACHVGGGTWTCTAAT-30) to the V3–V4 variable region of the 16S rRNA gene, while in the second stage, PCR amplification was performed on the first stage of PCR products using the Fluidigm Access Array barcoded primers. The 16S rRNA gene sequencing was performed using MiSeq according to the Illumina protocol. Based on the distribution of reads per barcode, the amplicons were pooled, re-pooled and purified. Then, the re-pooled libraries were loaded onto a Miniseq flow cell and sequenced (2×153 paired-end reads). Library preparation, pooling, and sequencing were performed at the Genome Research Core (GRC) within the Research Resources Center (RRC) at the University of Illinois Chicago.

#### Bioinformatic analysis

After DNA sequencing, all possible raw paired-end reads were evaluated and merged using the PEAR (Paired-End reAd mergeR) software (http://www.exelixis-lab.org/web/software/pear.) ^66^. To filter out low-quality reads, quality trimming was processed based on quality threshold (p = 0.01) and length parameters (minimum length = 225). The adapter/primer sequences were trimmed from the reads (fwd = GTGCCAGCMGCCGCGGTAA, and rev = GGACTACHVGGGTWTCTAAT) and ambiguous nucleotides (N=1) were trimmed from the ends and reads with internal ambiguous nucleotides were discarded. Chimeric sequences are artifacts of the PCR process and occur when portions of two separate amplicons fuse during the amplification process. Sequencing chimera filtering was carried out using a standard chimera checking program(the reference-based USEARCH (http://www.drive5.com/usearch) algorithm) ^67^ to identify and remove the chimeric sequences from the dataset as compared with a reference database (silva_132_16S.97)^68^. The sequences that passed both reference and algorithm methods are retained, and operational taxonomic units (OTUs) were selected based on a 97% similarity representing each OUT. Dereplication and sequences clustering via naïve read simplification (a.k.a. sub-OTU processing) techniques were used to reduce read count complexity. First, sequences were dereplicated; then minor sequences, those with counts below a given threshold (i.e., master cutoff of 10, and cluster threshold of 97% similarity) were clustered to major sequences. Then, OTUs were annotated using the USEARCH algorithm^5^ to compare with the reference database (silva _132_16S.97) ^6^ to identify references and taxonomic assignment from kingdom to species.

After removing the identical sequences with ≥97% identity of each other to make it nonredundant, the identified taxa have total counts of 4,369,887(Min: 38.0, Max: 71935.0, Median: 57916.000 and Mean: 5,5315.025 per sample). On the phylum level, Bacteroidetes (total counts: 2,766,568, 63.31%) and Firmicutes (total counts: 1,407,939,32.22%) accounted for most of the microbial population in all samples (95.53%) and Proteobacteria accounted for 1.92% (total reads: 84,079), while all other taxa were less than 1%.

Before normalization, data are first filtered to remove read counts associated with particular taxa (i.e., Chloroplast, mitochondria, Mitochondria, Chloroplast), then read counts were normalized as fraction of total sequence counts in each sample to obtain relative sequence abundance. The OTUs were removed from down-stream analysis if they were not present in all samples, had relative abundance less than 0.01%.

### Statistical Analysis

All data are expressed as the mean ± SD. All statistical tests were 2-sided. The *p* values < 0.05 were considered statistically significant. Before choosing an appropriate statistical test or model, the Shapiro-Wilk normality test for the distribution were performed for all the measurement variables. The differences between samples for two groups were analyzed using unpaired student’s t-test, Welch’s *t* test or Wilcoxon test based on the distribution of the testing variables. The differences between samples for more than two groups were analyzed using one-way ANOVA, two-way ANOVA or Kruskal-Wallis rank sum test as appropriate based on data distribution and the number of factors, respectively. The *p* values in ANOVA analyses and Kruskal-Wallis rank sum test were adjusted for correction of multiple comparisons using the Tukey or false discovery rate (FDR) method to ensure accurate results. Pairwise correlation analysis was conducted using Pearson or Spearman methods based on whether the testing variable was normally distributed or not. Regression plots in correlation analysis, line plots and scatter plots in longitudinal analysis were conducted using the ggplot2 package via R. To identify the core bacteria that were dynamically changed over time, we fitted two longitudinal models: Fast Zero-inflated Negative Binomial Mixed Modeling (FZINBMM) ^38^ and the linear mixed-effects models ^39, 40^. All the statistical analyses of microbiome data were performed using the R software (R version 4.0.4, 2021, The R Foundation for Statistical Computing Platform). Other statistical analyses were performed using GraphPad Prism 6 (GraphPad, Inc., San Diego, CA., USA).

## Author Contributions

YZ performed the cellular models, and animal studies, and the detailed analyses of the results; DO and SG, participated in animal studies; YX and YZ, perform the analyses of the microbiome and signaling pathways; YZ YX, and JS, prepared the figures and the draft text; YX contributed to the statistical analysis of data and the draft text; and JS obtained funds, designed the study, and directed the project. All authors contributed to the writing of the manuscript.

## Acknowledgements/Funds

We would like to acknowledge the VA Merit Award 1 I01BX004824-01, the NIDDK/National Institutes of Health grant R01 DK105118, and R01DK114126 to Jun Sun. The study sponsors play no role in the study design, data collection, analysis, and interpretation of data. The contents do not represent the views of the United States Department of Veterans Affairs or the United States Government.

