## Supplemental figure 1&2 for "Aberrant enteric neuromuscular system and dysbiosis in amyotrophic lateral sclerosis"

**Figure S1. Time-dependent pathophysiological changes in the colon of the SOD1$^{G93A}$ ALS mice.** ***(A)*** At the age of 2-month-old, intestinal mobility (whole gut transit time), started to increase while rotarod test time (Latency to fall time), forelimb grip strength and hindlimb grip strength started to decrease in age matched SOD1$^{G93A}$ mice compared to WT mice. (Data are expressed as mean ± SD. n = 6, two-way ANOVA test, *P < 0.05, ***P < 0.001). ***(B)*** At the age of 2-month-old, the expression of SMMHC protein started to decrease while the expression of GFAP protein, PGP9.5 protein and aggregation of human-SOD1$^{G93A}$ protein started to increase in age-matched SOD1$^{G93A}$ mice compared to WT mice (Data are expressed as mean ± SD. n = 6; the expression of SMMHC protein and PGP9.5 protein was analyzed by using two-way ANOVA test, while the expression of PGP9.5 protein and aggregation of human-SOD1$^{G93A}$ protein was analyzed by using Kruskal-Wallis test, **P < 0.01, ***P < 0.001).

### Fig. S1

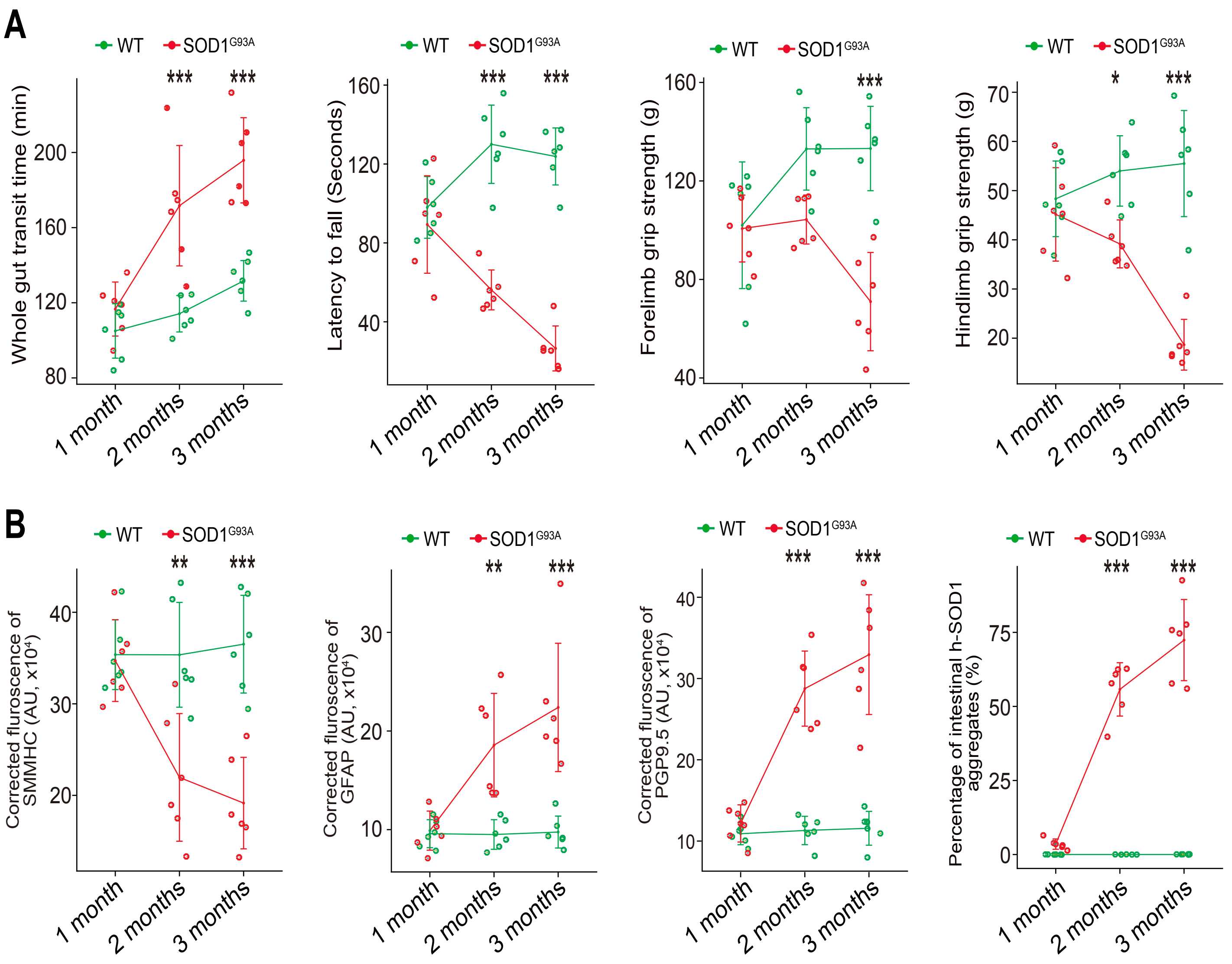

A
WT
SOD1G93A
Whole gut transit time (min)
Latency to fall (Seconds)
Forelimb grip strength (g)
Hindlimb grip strength (g)
1 month
2 months
3 months
B
Corrected fluroscence of SMMHC (AU, x10⁴)
Corrected fluroscence of GFAP (AU, x10⁴)
Corrected fluroscence of PGP9.5 (AU, x10⁴)
Percentage of intestinal h-SOD1 aggregates (%)

**Figure S2. Association of the altered ENS and increased aggregation of human-SOD1$^{G93A}$ in 3-month-old SOD1$^{G93A}$ mice in longitudinal studies.** ***(A)*** The correlation analyses between intestinal motility (Whole gut transit time) and other indexes i.e. latency to fall, forelimb grip strength, staining intensity of GFAP, aggregation of human SOD1$^{G93A}$ protein in 3-month-old SOD1$^{G93A}$ mice, compared with WT mice. (P values were labeled in figures, n = 6). Intestinal motility slow link with decreased forelimb grip strength; increased staining intensity of GFAP; increased aggregation of human SOD1$^{G93A}$ protein in 3-month-old SOD1$^{G93A}$ mice, compared with the WT mice. ***(B)*** The correlation analysis between aggregation of human SOD1$^{G93A}$ protein and other indexes, i.e. latency to fall, forelimb grip strength, staining intensity of SMMHC, staining intensity of GFAP protein in 3-month-old SOD1$^{G93A}$. (P values were labeled in figures, n = 6). Increased aggregation of human SOD1$^{G93A}$ protein links with decreased forelimb grip strength; decreased staining intensity of SMMHC; increased staining intensity of GFAP in the 3-month-old SOD1$^{G93A}$ mice, compared with the WT mice. ***(C)*** The correlation analysis between staining intensity of SMMHC and staining intensity of GFAP; staining intensity of PGP9.5 protein in the 3-month-old SOD1$^{G93A}$ mice, compared with the WT mice. (P values were labeled in figures, n = 6). Decreased staining intensity of SMMHC link with increased staining intensity of GFAP and PGP9.5 in 3-month-old SOD1$^{G93A}$ mice, compared with the WT mice.

### Fig. S2

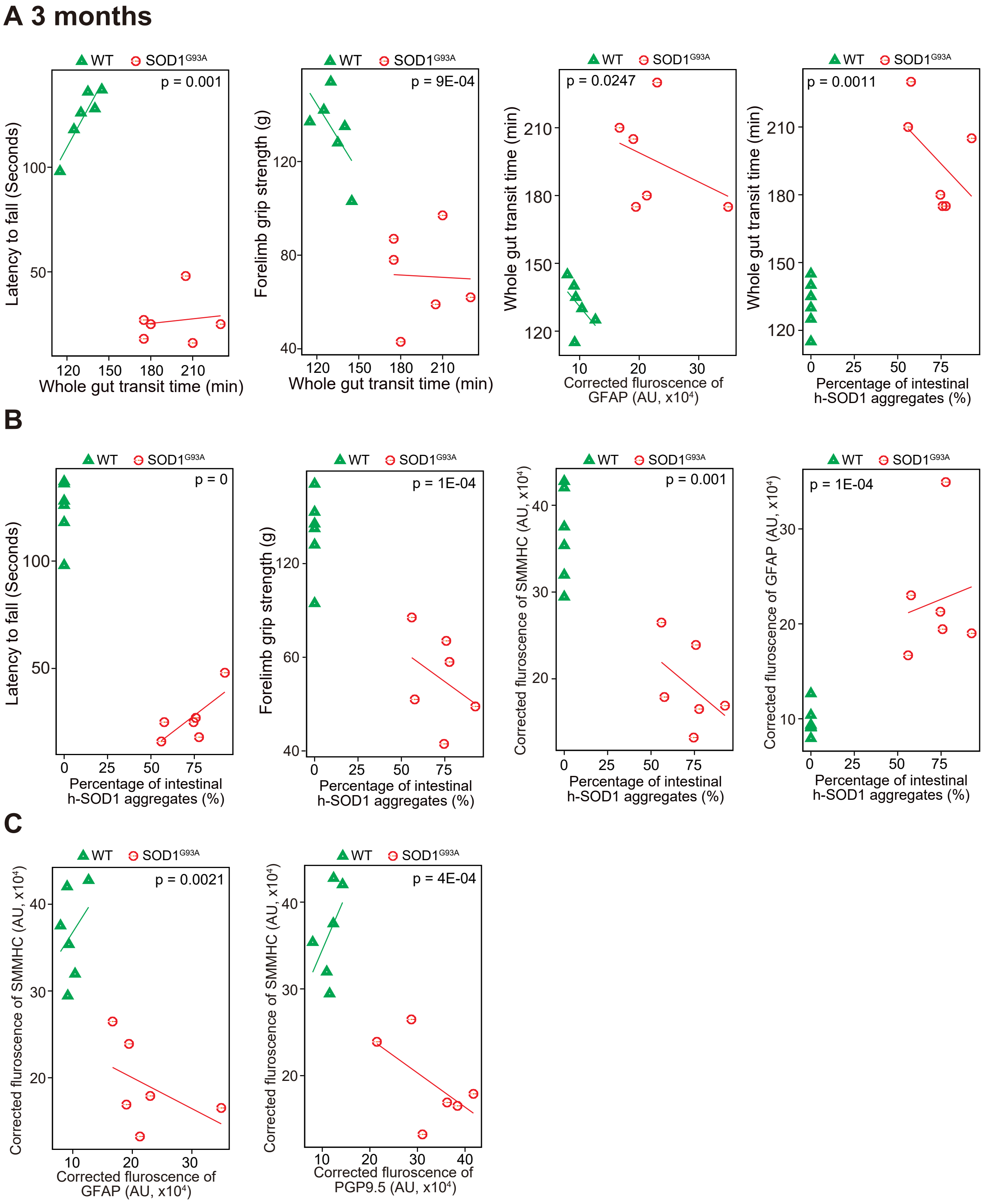
A 3 months
WT
SOD1G93A
p = 0.001
Latency to fall (Seconds)
Whole gut transit time (min)
p = 9E-04
Forelimb grip strength (g)
p = 0.0247
Whole gut transit time (min)
Corrected fluroscence of GFAP (AU, x10⁴)
p = 0.0011
Percentage of intestinal h-SOD1 aggregates (%)
B
p = 0
p = 1E-04
p = 0.001
Corrected fluroscence of SMMHC (AU, x10⁴)
Corrected fluroscence of GFAP (AU, x10⁴)
C
p = 0.0021
p = 4E-04
Corrected fluroscence of PGP9.5 (AU, x10⁴)
